# Prolonged exposure treatment following chronic social defeat stress in mice reduces social avoidance and stress responses

**DOI:** 10.1101/2025.04.28.651027

**Authors:** Jiwon Kim, Chung Sub Kim

## Abstract

**Background:** Chronic social defeat stress (CSDS) leads to persistent behavioral deficits, such as social avoidance and abnormal stress responses, including freezing. Despite these fear-based responses observed during and after social defeat, the effects of prolonged exposure (PE) treatment - a well-established therapy for post-traumatic stress disorder (PTSD)—have not been thoroughly investigated in this context.

**Methods:** Socially defeated male mice were subjected to 12 days of PE treatment. We determined the efficacy of PE treatment in physically defeated mice by measuring stress responses, including freezing, jump escape, and social interaction, before and after the intervention.

**Results:** After 12 days of PE treatment, male mice that had been physically defeated showed a marked decrease in freezing behavior. Furthermore, PE treatment changed the phenotype of mice from susceptible to resilient by reducing social avoidance during the social interaction test.

**Conclusions:** These results provide a new preclinical method for investigating behavioral recovery and adaptation by showing that PE treatment can reverse significant behavioral deficits caused by social defeat stress. A new framework for examining individual variability in therapeutic outcomes after chronic stress exposure is provided by the emergence of both treatment-responsive and treatment-resistant phenotypes.

## 1. Introduction

Chronic stress is a major risk factor for the onset of mental health disorders, such as depression and anxiety, which have prevalence rates of 4.73% and 6.16% in the United States, respectively (Dattani et al., 2021). While the majority of persons do not experience depression or anxiety disorders after stressful life events, a susceptible subgroup does, underscoring the need for sub-populational strategies to elucidate the underlying pathogenic pathways. Among stress-related conditions, posttraumatic stress disorder (PTSD) is a severe and long-lasting mental disorder that can develop after exposure to traumatic, life-threatening events. Similar to other stress-related disorders, PTSD occurs in only a subset of exposed individuals, while others demonstrate resilience. Moreover, severe physiological or psychological responses following a traumatic incident are associated with an increased probability of developing PTSD, since these initial reactions might strengthen traumatic memories and disrupt brain circuits responsible for stress management (McFarlane, 2010; Shin and Liberzon, 2010; Yehuda and LeDoux, 2007). The established association between trauma and PTSD is well-documented; however, symptoms may emerge months or even years after the traumatic event, complicating timely diagnosis and treatment.

Chronic social defeat stress (CSDS) is a prevalent mouse model used to investigate psychosocial stress, with the objective of clarifying the processes that contribute to behaviors like social avoidance and anhedonia (Kim et al., 2022a; Kim et al., 2025; Kim et al., 2022b; Krishnan et al., 2007). Conventional CSDS research mostly emphasizes post-defeat behaviors, often overlooking the rapid stress responses that transpire during the social defeat phase. The resident-intruder paradigm demonstrates that an intruder mouse is subjected to aggression from a resident mouse, which results in an intruder mouse engaging in social avoidance, abnormal fear memory, and impaired fear extinction that are comparable to that of bullied or defeated individuals (Golden et al., 2011; Kim et al., 2022a; Kim et al., 2025). The effects of social defeat are long-lasting, at least 3 months for socially defeated mice (Kim et al., 2022a). The CSDS method’s ability to create individual differences is a big plus because it lets researchers divide animals into two groups: those that are resilient and those that are not (Krishnan et al., 2007). The aforementioned variety bears a striking resemblance to the individual differences commonly observed in stress-related mental disorders (American_Psychiatric_Association, 2013). It is possible that prolonged exposure (PE) therapy could help these mice because they behave in ways that are similar to those seen in people with PTSD, like avoiding social situations (e.g., social avoidance), not enjoying things (e.g., anhedonia), and having trouble remembering where they put things (e.g., spatial working memory)(Kim et al., 2022a; Krishnan et al., 2007).

In the treatment of PTSD, the cognitive-behavioral intervention known as PE therapy is aimed to promote the progressive confrontation and processing of trauma-related memories, emotions, and avoidance behaviors. This therapy has been empirically validated. (McLean and Foa, 2011; Powers et al., 2010). Given that physically socially defeated mice exhibit fear-based freezing and social avoidance during the social defeat session and the social interaction test (Kim et al., 2025; Kim et al., 2022b), treating these behavioral deficits with prolonged exposure treatment opens a new avenue for investigating the mechanisms underlying PE treatment.

## 2. Methods

### 2.1 Animals

Male C57BL/6J mice (7–8 weeks old) were used in this study. To select aggressors, 3-to 6-month-old male CD-1 mice were utilized, while 7-to 8-week-old male CD-1 mice were chosen as non-aggressors. Ovariectomized 2-to 3-month-old female CD-1 mice served as partners for male CD-1 mice, and 5-week-old male CD-1 mice underwent surgical castration. Before undergoing chronic social defeat stress and prolonged exposure treatment, animals were housed in groups of 2 to 5 per cage under a 12-hour light/dark cycle (lights on at 6 am and off at 6 pm) with ad libitum access to food and water. Housing conditions for each behavioral experiment are described in their respective sections. All procedures involving animals were approved by the Institutional Animal Care and Use Committee (IACUC) of Augusta University.

### 2.2 Ovariectomy

The surgical procedure for ovariectomy was performed as previously described(Kim et al., 2025; Kim et al., 2022b). Carprofen (5 mg/kg, s.c., Covetrus) was administered preoperatively. Female mice were anesthetized with 2% isoflurane (0.5 L/min oxygen) in an induction chamber for two minutes before being positioned ventrally. Their fur was clipped around the third lumbar vertebra and the surgical site disinfected. Two 0.5 cm dorsal incisions were made just caudal to the last rib, and blunt forceps were used to separate tissue layers. A small peritoneal incision (<0.5 cm) was made on each side, and the ovarian fat pad was retracted to expose the oviduct. The ovary was ligated (∼0.5 cm from the ovary), severed with sterile micro scissors, and the remaining oviduct reinserted. The muscle layer and skin were closed with absorbable sutures. Mice were placed in a recovery cage on a heated surface and monitored every 15 minutes until full recovery before being returned to their housing room.

### 2.3 Surgical Castration

The surgical procedure for castration was performed as previously described(Kim et al., 2025; Lofgren et al., 2012). Castrations were performed to prepare socially defeated mice for prolonged exposure treatment. All surgical instruments were autoclaved before the procedure. Carprofen (5 mg/kg, s.c., Covetrus) was administered preoperatively. Five-week-old male CD-1 mice were anesthetized with 2% isoflurane (0.5 L/min oxygen) throughout the procedure. The inguinal-perineal region was clipped and disinfected with three alternating wipes of alcohol and betadine. A scrotal incision was made, and the skin and body wall caudal to the penis were opened with sterile scissors. The testicles were identified and retracted, and the testicular artery and vein were double-ligated and severed. The body wall and skin were closed with absorbable sutures. Mice were placed in a recovery cage on a heated surface and monitored every 15 minutes until fully recovered, after which they were returned to their housing room.

### 2.4 Selection of aggressors

The selection of aggressors was conducted as previously described (Kim et al., 2022a; Kim et al., 2025; Kim et al., 2022b). Male CD-1 mice, aged 3 to 6 months, were co-housed with ovariectomized female CD-1 mice in large cages for at least two weeks. On the screening day, the female mouse was removed, leaving the male CD-1 mouse in the cage. A ‘screener’ C57BL/6J male mouse (7 weeks old) was then introduced into the cage for a 3-minute session, during which the latency of the CD-1 mouse to attack the screener was recorded. After each session, the screener mouse was removed, and the female CD-1 mouse was returned to co-house with her partner. This screening procedure was repeated for three consecutive days. Male CD-1 mice with an attack latency of less than 60 seconds in at least two sessions were selected as aggressors for the social defeat experiments. The selection of aggressors was performed between 4 pm and 6 pm.

### 2.5 Selection of nonaggressors

The selection of nonaggressors was conducted as previously described(Kim et al., 2025). Sexually inexperienced male CD-1 mice and castrated male CD-1 mice (castrated at 5 weeks old) were screened as non-aggressors at 7–8 weeks of age. On the screening day, a male CD-1 mouse and a 7-week-old “screener” C57BL/6J mouse were placed in a large new cage for a 10-minute session, during which any aggressive behaviors (e.g., attack bites) by the CD-1 mouse were recorded. Following each session, the mice were returned to their individual home cages. The screening method was conducted on three successive days. CD-1 mice who shown no aggressive behavior throughout all three sessions were designated as non-aggressors for the prolonged exposure treatment trial. Screening was conducted between 4 pm and 6 pm.

### 2.6 Chronic physical social defeat stress

Social defeat was generated using a resident–intruder paradigm as previously described (Kim et al., 2022a; Kim et al., 2025; Kim et al., 2022b). For the chronic physical social defeat procedure, male C57BL/6J experimental mouse was introduced into the large home cage of an aggressor CD-1 mouse for a 10-minute session. After the session, the physically defeated mouse was co-housed with the CD-1 aggressor but separated by a perforated Plexiglass divider, allowing only sensory contact for the next 24 hours. The following day, the defeated mouse was re-exposed to a new CD-1 aggressor for another 10-minute session, followed by 24 hours of sensory contact. This procedure was repeated daily for 12 days. Control C57BL/6J mice were housed in pairs in cages identical to those used for socially defeated mice. After the final social defeat stress session, both control and defeated mice were individually housed in standard cages. The social defeat stress procedure was performed between 4 pm and 6 pm.

### 2.7 Social interaction test

Social interaction test was performed as previously described (Kim et al., 2022a; Kim et al., 2025; Kim et al., 2022b). The test was conducted 48 hours after the final social defeat stress session. Mice were acclimated to a behavior room under red-light conditions (∼4 lux) for at least 1 hour, 2 days prior to the test. The apparatus consisted of a box (18 × 18 × 12 inches) with a perforated Plexiglass cage (4 × 4 × 12 inches) positioned along one side. The social interaction test followed a two-trial procedure under red-light conditions (∼4 lux). In the first session, the experimental mice were placed in the corner of the open arena and allowed to explore freely for 2.5 minutes without a social target. In the second session, an unfamiliar CD-1 male aggressor was introduced into the perforated Plexiglass cage as a social stimulus, and the experimental mice were again allowed to explore for 2.5 minutes. The surface of the open field was cleaned with 70% ethanol to remove residual odors after each trial. A CCD camera positioned above the apparatus recorded the behavior. The social interaction ratio was calculated as the ratio of the time spent in the interaction zone in the presence of a social target to the time spent in the interaction zone without a social target. Mice with scores greater than 1 were defined as “resilient-like phenotype,” while those with scores less than 1 were defined as “susceptible-like phenotype”(Golden et al., 2011). A custom-written program was used to analyze the behavioral data. The behavioral experiment was conducted under blinded conditions between 4 pm and 6 pm.

### 2.8 Stress responses during the social defeat session

Behavioral stress responses, including freezing, social approach, and socialization, were measured during a 10-minute social defeat session under blinded conditions. Freezing is defined as a defensive response to fear, exemplified by the presence of a CD-1 mouse, during which the mouse remains motionless to avoid detection. Social approach is defined as the defeated mouse actively moving toward and engaging with a CD1 non-aggressor. Socialization is defined as the defeated mouse remaining in close proximity to a CD-1 aggressor without showing avoidance behavior.

### 2.9 Prolonged exposure treatment

Prolonged exposure treatment was performed as previously described(Kim et al., 2025). Pre-selected non-aggressors at 7-8 weeks of age were used. Sexually inexperienced male CD-1 mice and castrated male CD-1 mice (castrated at 5 weeks old) were used. On the screening day, a male CD-1 mouse and a 7-week-old “screener” C57BL/6J mouse were placed in a large new cage for a 10-minute session, during which any aggressive behaviors (e.g., attack bites) by the CD-1 mouse were recorded. Following each session, the mice were returned to their individual home cages. The screening method was conducted on three successive days. CD-1 mice who shown no aggressive behavior throughout all three sessions were designated as non-aggressors for the prolonged exposure treatment trial. PE treatment was conducted between 4 pm and 6 pm.

### 2.10 Statistical Analysis

Statistical comparisons were performed using a simple linear regression test, paired t-test (Wilcoxon signed-rank test), or unpaired t-test (Mann-Whitney U test) using GraphPad software. Statistical significance was considered at ^*^P < 0.05.

## 3. Results

### 2.1 Selection of aggressors

Since the chronic social defeat stress (CSDS) paradigm heavily relies on the aggressive behaviors of resident CD-1 mice, such as attack bites, we conducted a three-day selection process to identify suitable aggressors (Fig. 1A). To achieve this, 7-to 8-week-old male screener mice were subjected to the resident-intruder paradigm. Out of 93 CD-1 male mice screened, 29 exhibited strong aggressive behavior, as indicated by an attack latency of less than 60 seconds (Fig. 1B, 1C). The observed strong negative correlation between attack latency and the number of attack bites suggests that a shorter attack latency is a reliable indicator of aggressive behavior (Fig. 1D).

**Fig. 1.**
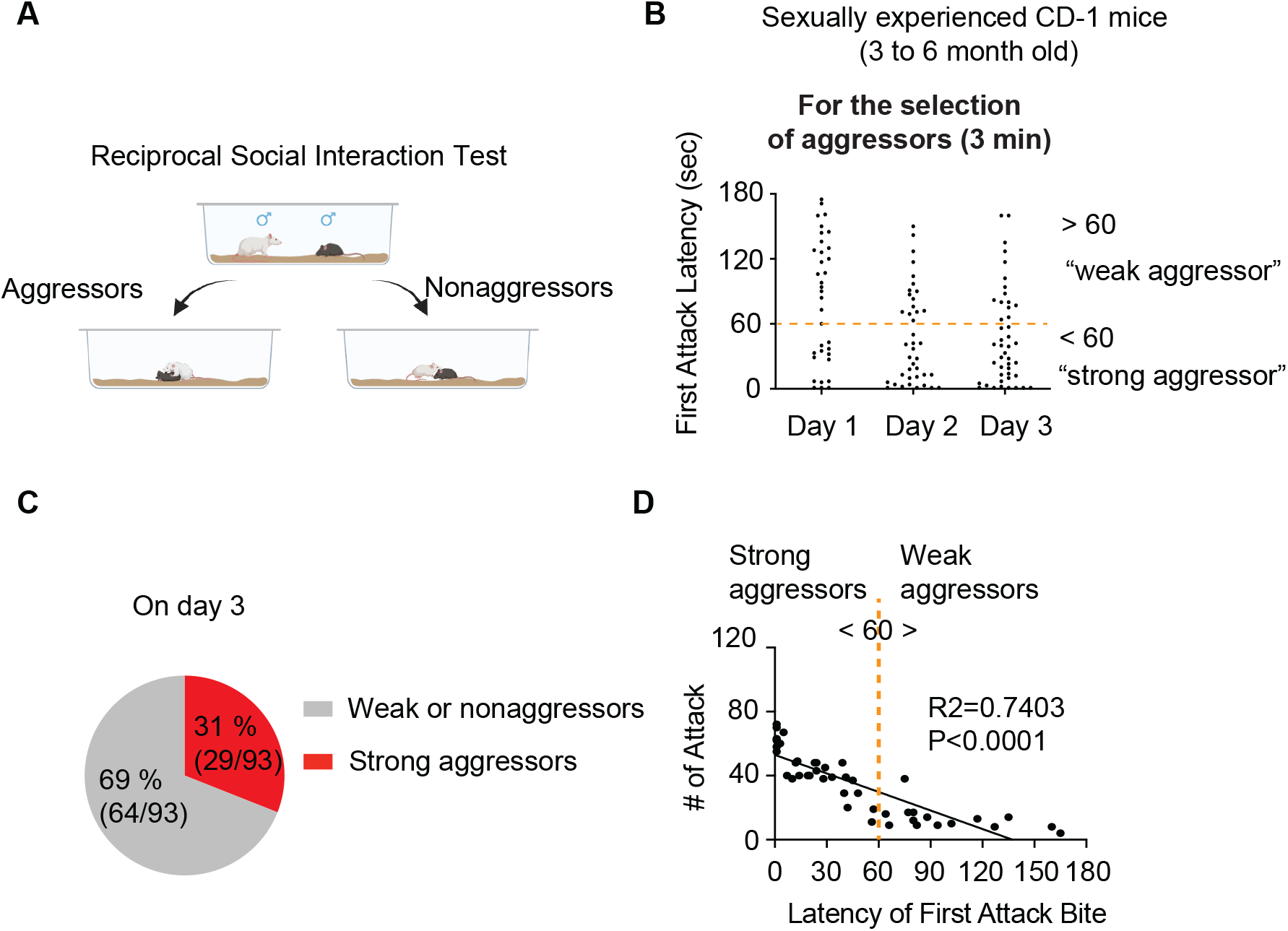
Selection of aggressors. **(A)** Illustration showing the reciprocal social interaction test used to screen aggressors. **(B)** The latency of the first attack bite during aggressor selection. **(C)** A pie chart depicting the percentages of strong aggressors versus weak or non-aggressors on the third day of selection. **(D)** On day 3, attack bites and attack latency show an inverse correlation. A simple linear regression analysis was conducted to examine the relationship between the number of attack and attack latency. Panel **(A)** was created with BioRender.com.

### 3.2 Selection of nonaggressors

We selected non-aggressive CD-1 mice by using a combination of sexually inexperienced, castrated CD-1 males (castrated at 5 weeks of age) and sexually inexperienced young CD-1 males (7-8 weeks old). Of 34 sexually inexperienced young CD-1 mice, 18 showed non-aggressive behavior throughout the 10-min screening across 3 consecutive days (Fig. 2A, 2B); out of 10 castrated CD-1 mice, 9 showed non-aggressive behavior for 3 consecutive days (Fig. 2C, 2D). 27 of 44 CD-1 mice overall fit our criteria for inclusion into the 12-day PE treatment (Fig. 2E, 2F).

**Fig. 2.**
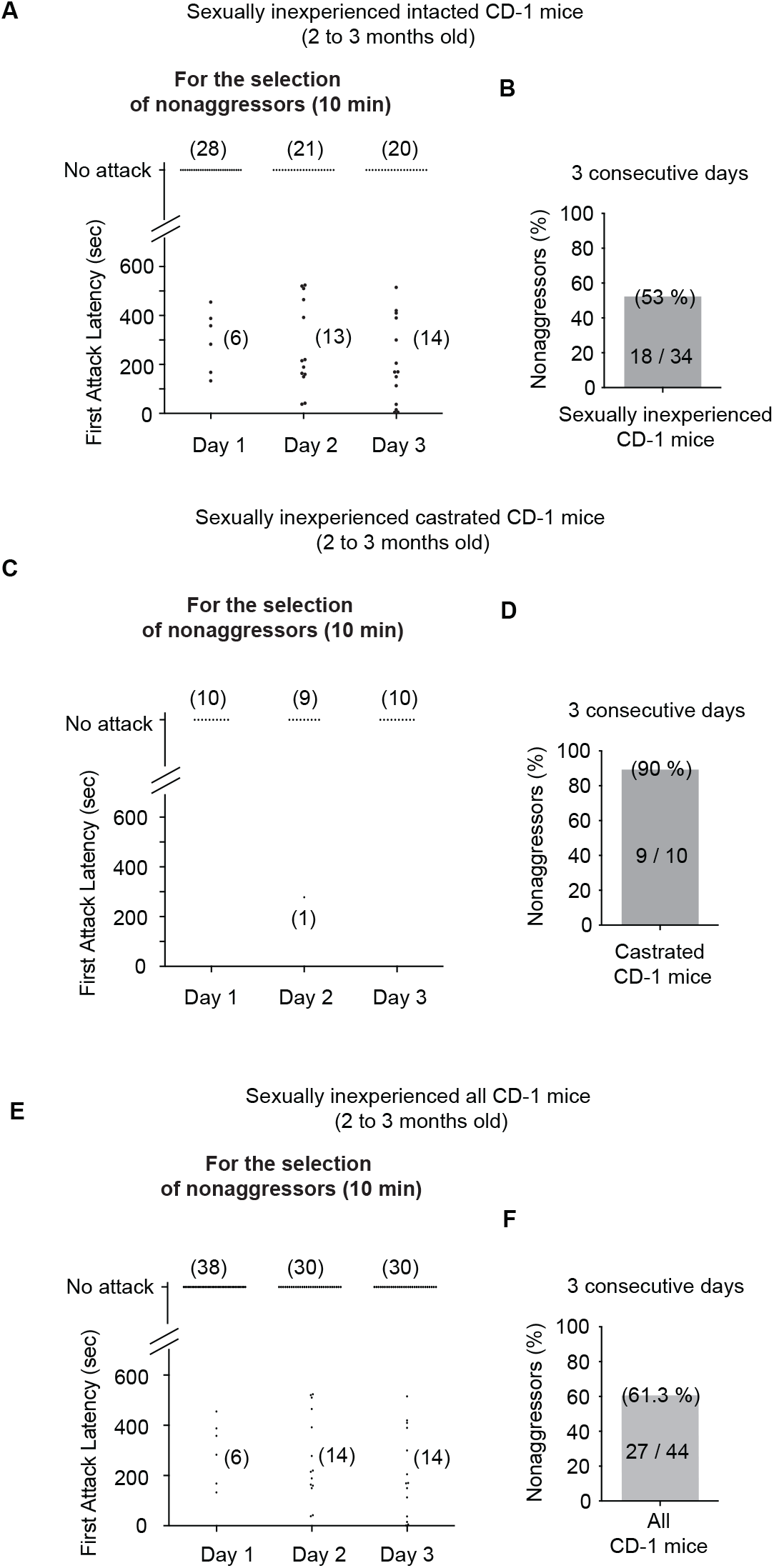
Selection of non-aggressors. **(A)** Latency of the first attack bite during non-aggressor selection in sexually inexperienced male CD-1 mice. **(B)** The percentages of non-aggressors over three consecutive days. **(C)** Latency of the first attack bite during non-aggressor selection in sexually inexperienced castrated CD-1 mice. **(D)** The percentages of non-aggressors over three consecutive days. **(E)** Combined latency of the first attack bite during non-aggressor selection in both sexually inexperienced intact and castrated male CD-1 mice. **(F)** The percentages of non-aggressors over three consecutive days.

### 3.3. Treatment with prolonged exposure improves behavioral abnormalities caused by social defeat

Male mice aged 7 to 8 weeks were subjected to CSDS. CSDS-treated mice spent significantly less time in the interaction zone when a novel aggressor was present (Fig. 3C), while no difference was observed in the absence of a social target (Fig. 3B). Social interaction ratios were significantly decreased in the CSDS-treated group (Fig. 3D), with 5 out of 12 mice displaying a resilient-like phenotype and 7 out of 12 mice displaying a susceptible-like phenotype (Fig. 3E). We then conducted 12 days of prolonged exposure (PE) treatment for socially defeated mice. According to earlier research, mice exposed to 6 days of SDS showed no social deficits (such as social avoidance and anxiety) after just 6 days of non-social defeat exposure (Kim et al., 2022b). Based on this, a minimum of 6 days of PE treatment might be necessary for recovery in mice subjected to a longer SDS duration. However, to better match the exposure duration of 12 days of SDS, we extended the PE treatment to 12 days, ensuring a balanced and systematic approach. After the social interaction test, each CSDS-treated mouse was housed separately and stayed that way for the PE treatment period. Because 12 days of social isolation in control mice significantly affected social behavior during the second social interaction test, the control group was not included in this test. The PE-treated mice were re-exposed to the social interaction test both with and without a new aggressor. Interestingly, PE-treated mice spent more time in the interaction zone when a new aggressor was present (Fig. 3G) although no effect was seen in the absence of a social target (Fig. 3F). Social interaction ratios (Fig. 3H) showed significant increase as a result. Especially, 4 of 12 mice displayed a PE-resistant-like phenotype and 8 of 12 mice showed a PE-responsive-like phenotype (Fig. 3I). During PE treatment (Fig. 3J), we observed a significant reduction in freezing behavior on Day 12 compared to Day 1 (Fig. 3K). Additionally, the number of social approaches (Fig. 3L) and the duration of socialization (Fig. 3M) were significantly increased on Day 12 compared to Day 1.

**Fig. 3.**
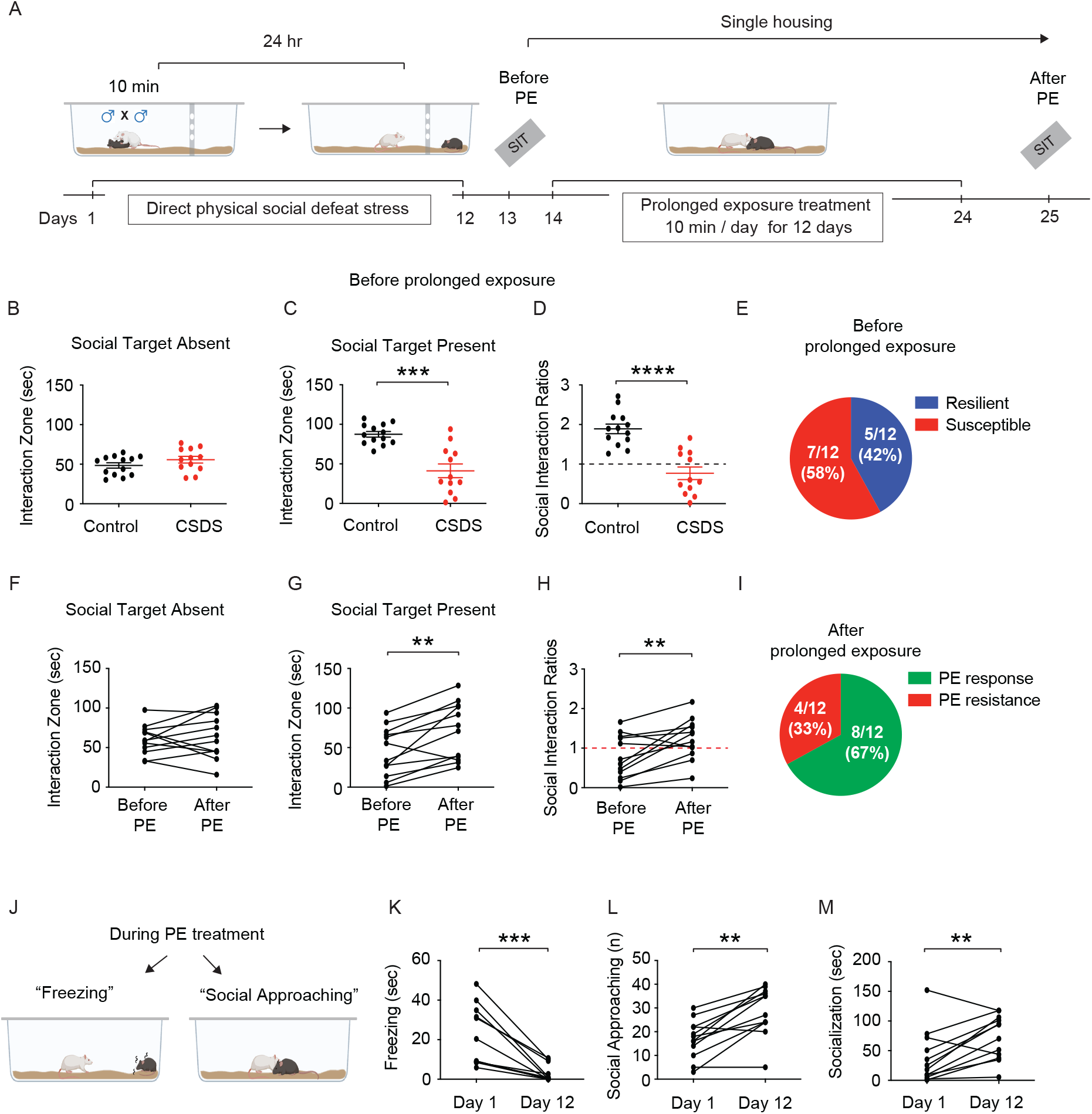
Prolonged exposure treatment reduced social defeat-induced behavioral deficits. (**A**) Schematic diagram of the experimental design for chronic physical social defeat stress, prolonged exposure treatment, and social interaction test. (**B**) Time spent in the interaction zone in the absence of a social target following the 12-day CSDS. (**C**) Time spent in the interaction zone in the presence of a social target (i.e., a novel aggressor) following the12-day CSDS. A Mann-Whitney test was performed. (**D**) Social interaction ratios. The Mann-Whitney test was performed. (**E**) Proportion of resilient- and susceptible-like phenotypes following the 12-day CSDS. (**F**) Time spent in the interaction zone in the absence of a social target before and after the 12-day prolonged exposure treatment. (**G**) Time spent in the interaction zone in the presence of a social target before and after the 12-day prolonged exposure treatment. A Wilcoxon matched-pairs signed rank test was performed. (**H**) Social interaction ratios before and after the 12-day prolonged exposure treatment. The Wilcoxon matched-pairs signed rank test was performed. (**I**) Proportion of PE-responsive- and PE-resistant-like phenotypes after prolonged exposure treatment. (**J**) Illustration of freezing and social approaching behaviors during prolonged exposure treatment. (**K**) Freezing time during prolonged exposure treatment on Day 1 and Day 12. The Wilcoxon matched-pairs signed rank test was performed. (**L**) The number of social approaching during prolonged exposure treatment on Day 1 and Day 12. The Wilcoxon matched-pairs signed rank test was performed. (**M**) Socialization time during prolonged exposure treatment on Day 1 and Day 12. The Wilcoxon matched-pairs signed rank test was performed. Data are expressed as mean ± SEM. **P<0.01, ***P<0.001, and ****P<0.001.

## 4. Discussion

This work highlights a unique strategy with significant translational value since it is the first case of PE therapy being used as a treatment for physically socially defeated mice. Similar to human prolonged exposure therapy (McLean and Foa, 2011), which uses non-threatening items or cues associated with traumatic events, selecting non-aggressor mice is crucial. This is because physically, socially defeated mice specifically exhibit fear-based social avoidance and freezing behaviors in response to aggressors (Kim et al., 2022a; Kim et al., 2025; Kim et al., 2022b). Aggressive behavior of CD-1 mice against C57/BL6J mice in a new cage setting seems to be influenced by physical size and previous sexual experience (Barr, 1981; Kim et al., 2022b; Schuett, 1997). Presumably because of their apparent dominance advantage, larger CD-1 mice are more likely to show hostility against smaller C57 males. Furthermore, sexually inexperienced CD-1 males show less inclination for violence as sexual experience is known to increase territoriality and dominance-related activities in rats (Barr, 1981). These results imply that aggressive interactions between male mice in new social environments are much influenced by physical and sensory aspects. In a previous study, selected male aggressors housed in a single cage initially showed strong aggressive behavior during the first 6 days of CSDS but lost their interest in attacking intruders for the rest of the 6 days of CSDS, resulting in physically, socially defeated mice showing normal-like behaviors in the social interaction test and open field test (Kim et al., 2022b). This result suggests that exposure therapy could potentially address anomalous social conduct resulting from direct social defeat experiences. We therefore selected nonaggressors with sexually inexperienced young CD-1 mice and sexually inexperienced and castrated young CD-1 mice. Unlike selecting aggressors, screener mice (e.g., C57BL/6J) were exposed to similar size of CD-1 mice for 10 min over 3 consecutive days. We only selected mice showing non-aggressive behaviors such as attack bite during free social behavioral test in a large new mouse cage for 3 consecutive days. Notably, 9 out of 10 surgically castrated young CD-1 mice showed non-aggressive behaviors (Fig 2D), indicating sexual experience or male hormone may play a role in male aggression.

PE therapy is a well-researched cognitive-behavioral method used in therapeutic settings to lessen symptoms of stress-related disorders, including PTSD, by promoting the gradual confrontation and processing of trauma-related memories and emotions. Indeed, prolonged exposure treatment to witness-expose male mice effectively improves freezing and social behaviors during the witnessing social defeat sessions (Day 1 vs. Day 12 of PE treatment), resulting in a shift of some susceptible phenotypes to resilient-like phenotypes (i.e., PE-responsive mice)(Kim et al., 2025). However, it has not yet been reported whether physically defeated mice have been treated with PE treatment. While our recently accepted study demonstrated the efficacy of PE treatment specifically in witness-exposed mice (Kim et al., 2025), the effects of this intervention on physically defeated mice were not examined. To address this critical gap, the present study investigates whether PE treatment can also improve stress-related behaviors in physically defeated mice, a population that more closely models direct trauma exposure. By using PE treatment in a preclinical model of physically defeated mice, we extend the treatment paradigm often used in human patients to an animal model that mimics significant behavioral symptoms, such as social avoidance and decreased stress regulation. It is important to note that although a significant decrease in freezing behavior and an increase in social behavior were observed on the final day of PE treatment (Fig. 3), a subset of mice exhibited a PE-resistant phenotype (Fig. 3), which is similar to the variability that is seen in the response to PE therapy in human patients suffering from PTSD (Bradley et al., 2005; Bryant, 2019).

Considering that pharmacological therapies, such as ketamine treatment, have demonstrated effectiveness in cases of treatment-resistant depression or PTSD (Philipp-Muller et al., 2021; Shiroma et al., 2024; Shiroma et al., 2020), it would be beneficial to test ketamine within this framework in order to evaluate whether or not mice who are resistant to PE may be changed towards a phenotype that is sensitive to PE. Following PE treatment, we observed a significant change in social defeat-induced stress responses (Fig. 3K-3M). Freezing behavior was significantly reduced, and self-motivated socialization was significantly increased on day 12 of PE treatment compared to on day 1 of PE treatment (Fig. 3K). Consistent with this observation, fear-based social avoidance induced by social defeat was significantly improved following PE treatment (Fig. 3H). Notably, there were PE-resistant and PE-responsive phenotypes following PE treatment (Fig. 3I) consistent with the variability of human PTSD with PE therapy (Bradley et al., 2005; Bryant, 2019). These results suggested that abnormal stress responses and impaired social behavior could be treated by PE treatment, and it may provide the novel approach to investigate the mechanisms underlying the individual differences in PE treatment.

## 5. Conclusion

This work introduces an innovative method with some major improvements to our preclinical model stress assessment strategy. One of the key ideas here is the emphasis on monitoring stress reactions throughout social defeat session, which might rather assist in predicting animal social behavior later. This paper introduces the use of prolonged exposure treatment to physically defeated mice, a novel approach that has not been previously undertaken. This method provides an innovative means to investigate possible interventions for stress-related behaviors. These novel methodologies enhance the model’s applicability for examining psychiatric disorders associated with persistent stress. The findings indicate potential for comprehending and addressing individual variations in animal responses to stress and therapy. Employing PE therapy within this framework also paves the way for future study, perhaps resulting in improved treatment alternatives for conditions such as PTSD.

## CRediT authorship contribution statement

**Chung Sub Kim:** Writing - original draft, review & editing, Investigation, Conceptualization. **Jiwon Kim**: Investigation.

## Acknowledgement

We thank Dr. Payne Y. Chang for providing the behavioral software. This work was supported by a start-up fund (to CSK) from the Medical College of Georgia at Augusta University, and by NIH grant R01MH134958 (to CSK).

## Data Availability

Data will be made available on request.

